# Anti-gene oligonucleotides targeting Friedreich’s ataxia expanded GAA•TTC repeats increase Frataxin expression

**DOI:** 10.1101/2024.01.25.577034

**Authors:** Negin Mozafari, Salomé Milagres, S. J. Rocha, Claudia Marina Vargiu, Fiona Freyberger, Osama Saher, Tea Umek, Pontus Blomberg, Per T. Jørgensen, C. I. Edvard Smith, Jesper Wengel, Rula Zain

## Abstract

Friedreich’s ataxia (FRDA) is a progressive, autosomal recessive ataxia caused, in the majority of cases, by homozygous expansion of GAA•TTC triplet-repeats in the first intron of the *frataxin* (*FXN*) gene. GAA•TTC repeat expansion results in the formation of non-B DNA intramolecular triplex structure (H-DNA) as well as changes in the epigenetic landscape at *FXN* loci and heterochromatin formation. Expansion of intronic GAA•TTC repeats is associated with reduced levels of *FXN* mRNA and protein resulting in disease development. Previously, we reported that DNA-binding anti-gene oligonucleotides (AGOs) targeting the GAA•TTC repeat expansion abolished H-DNA formation. Here, we demonstrate that targeting repeat-expanded chromosomal DNA using single-strand locked nucleic acid (LNA)-DNA mixmer AGOs increases FXN mRNA and protein expression in patient-derived cells. We examined numerous LNA-DNA AGOs and found that the design, length and their LNA composition have a high impact on the effectiveness of the treatment. Collectively, our results demonstrate the unique capability of specifically designed ONs targeting the GAA•TTC DNA repeats to upregulate *FXN* gene expression.

## INTRODUCTION

Friedreich’s ataxia (FRDA) was first described by Nikolaus Friedreich in 1877 ^1^. The clinical features of FRDA are ataxia, scoliosis, foot deformity, cardiomyopathy, and diabetes mellitus ^2,3^. The most affected tissues within the central nervous system (CNS) are the spinal cord and dorsal root ganglia^4^. Additionally, the peripheral nervous system, heart muscle, skeleton, and pancreas, are also affected ^5^. The patients normally begin to show symptoms during childhood and the first years of adolescence and lose their walking ability on average 10 to 15 years after disease onset ^6,7^.

FRDA is an inherited autosomal recessive ataxia with a prevalence of about 1:50,000 individuals. Homozygous expansion of the GAA•TTC triplet-repeat in intron 1 of the *frataxin* (*FXN)* gene is associated with FRDA in 98% of affected individuals. The remaining cases are related to heterozygosity, with GAA expansion on one allele and a loss-of-function variation, such as point mutations or deletions on the other ^8–10^.

*FXN* is a nuclear gene which encodes the FXN protein that localizes mainly into mitochondria; but can after maturation also be located in nuclei, endoplasmic reticulum and microsomes ^11,12^. FXN protein is responsible for the iron-sulphur (Fe-S) cluster biosynthesis acting as an allosteric activator ^13^. The role of Fe-S clusters in cells is diverse, from electron transfer and Fe regulation to DNA repair. Malfunctioning of these processes causes Fe-S deficiency and accumulation of toxic Fe in mitochondria^14^. The expansion of GAA•TTC repeats in the *FXN* gene results in reduced transcription and subsequently lowered levels of FXN protein, leading to mitochondrial Fe accumulation and increased cellular oxidative stress ^14–16^.

In FRDA, the length of the GAA•TTC repeats is associated with disease age-at-onset and severity. Pathogenic expanded alleles carry around 66-1700 GAA•TTC repeats, whereas a healthy allele contains 7 to 22 ^17,18^. While considered recessive, individuals carrying an expanded GAA•TTC repeat in only a single chromosome and no pathogenic expansions in the other allele (haploinsufficiency) may develop mild clinical features ^10,19,20^. Germline instability of expanded repeats is reported in both paternal and maternal transmission, with more frequent contractions when inherited from the paternal allele, while both expansions and contractions are derived from the maternal line ^21^. In premutation alleles harboring 22-60 repeats, the incidence of repeat instability increases ^22–24^. Apart from the intergenerational unstable transmission, the expansion of GAA•TTC repeats vary extensively within the individual’s tissues ^25,26^. Somatic instability of pathogenic expanded alleles is progressive throughout the lifetime ^27^, affecting tissues like the heart, dorsal root ganglia, cerebellum, pancreas, and spinal cord^25,26^.

Several factors are reported to cause disturbed transcription initiation or elongation thereby contributing to *FXN* gene silencing in FRDA ^18,28–32^. It has been suggested that expanded GAA•TTC repeats form non-B DNA structures like intramolecular triplex structure (H-DNA) or DNA-RNA structure (R-loop), reducing *FXN* mRNA and hence also protein levels. ^18,28,29^. Furthermore, epigenetic changes and heterochromatin formation are linked to *FXN* gene silencing at the expanded locus ^30^.

Currently, there is no cure available for FRDA, and the existing therapies only treat symptoms ^6,33^. In February 2023, the FDA approved Omaveloxolone (Skyclarys^TM^), making it the first and only approved drug for FRDA. Omaveloxolone activates nuclear factor erythroid 2-related factor 2 (Nrf2), a transcription factor suppressed in FRDA, and that is responsible for maintaining redox homeostasis and counteracting the production of reactive oxygen species. Nrf2 activation through Omaveloxolone has shown to improve mitochondrial function and relief of some symptoms ^34^. Additional disease-modifying trials are ongoing and comprise a variety of substances targeting pathways underlying FRDA pathogenesis ^11^. It has been reported that histone deacetylase inhibitors (HDACi) can restore FXN expression by increasing histone acetylation and subsequently activation of transcription *in vitro* and *in vivo* ^35^. Thus, derivatives of 2-aminobenzamide ^36,37^ and nicotinamide ^38^ partially upregulate FXN expression in both patient-derived cells and patients. Moreover, increased FXN mRNA and protein expression have been achieved by targeting the proposed R-loop with antisense oligonucleotides (ASOs) and duplex RNAs ^39,40^. Nevertheless, the effect of these ASOs and duplex RNAs in cell cultures were not translatable in an FRDA mouse model ^41^. A recent study focusing on gene therapy such as gene editing and gene replacement strategies using CRISPR-Cas9 technology, accomplished *FXN* upregulation in an FRDA mouse model ^42^. In an FRDA mouse model, intravenous delivery of a human *FXN* gene by an adeno-associated virus reversed the cardiomyopathy phenotype ^43^. Most recently, ASOs directed against the 5′ and 3′ untranslated region (UTR) of *FXN* mRNA were shown to increase FXN mRNA and protein expression in FRDA patient-derived cell lines ^44^.

Our approach is to address the root of the disease, which is the DNA repeat expansion. To this end, we have previously designed repeat-targeting AGOs to disrupt the H-DNA structure formed at GAA•TTC expanded repeats in plasmids in a sequence- and structure-specific manner ^45,46^. More recently we showed that our AGO modality, targeting the H-DNA at the GAA•TTC repeat expansion, prevented repeat expansion in a mammalian reporter model ^47^. Here we aim to activate *FXN* expression in FRDA patient-derived primary fibroblasts using GAA Locked nucleic acid/ DNA (LNA/DNA) mixmers with a fully phosphorothioate (PS) -modified backbone targeting the H-DNA structure in intron 1. The AGOs were chemically modified comprising of a PS ^48^ backbone to enhance resistance toward nucleases in mixmers containing LNA ^49^ to increase affinity toward the H-DNA forming GAA•TTC repeats. To our knowledge, this is the first report demonstrating activation of FXN expression by DNA repeat targeting AGOs. The GAA AGOs are reverse complementary to the template strand and significantly upregulated FXN mRNA and protein expression in a dose-dependent way. In contrast, the cognate CTT ONs are reverse complementary to the coding strand and pre-mRNA ^29^. and reduced FXN levels. We also evaluated the design of LNA/DNA mixmers and the LNA content, which shows that these are key to achieving efficient FXN upregulation.

## RESULTS

### GAA AGOs significantly enhance *FXN* mRNA expression

We have previously shown that modified AGOs, which bind to the GAA•TTC repeat expansions, resulted in different DNA complexes depending on their sequence. A GAA ON significantly disrupted the H-DNA triplex structure in plasmids containing pathogenic expanded (GAA•TTC)_115_ repeats while, conversely, a CTT ON enhanced triplex formation ^45,46^. We hypothesized that a GAA AGO could, by abolishing the H-DNA, facilitate *FXN* transcription and restore mRNA and protein levels. To validate this hypothesis, GAA AGOs were designed as LNA/DNA mixmers with a fully PS-modified backbone and directed against the pyrimidine motif triplex (Table 1 and Figure 1A). First, we designed and synthesized various GAA AGO mixmers of different lengths and LNA content. The GAA_15_ AGO contains 40% LNA; the first guanine and every second adenosine are LNA. Other 15 and 16 mer GAA AGOs with different LNA patterns were also synthesized (Table 1 and Figure 1A).

**Table 1:**
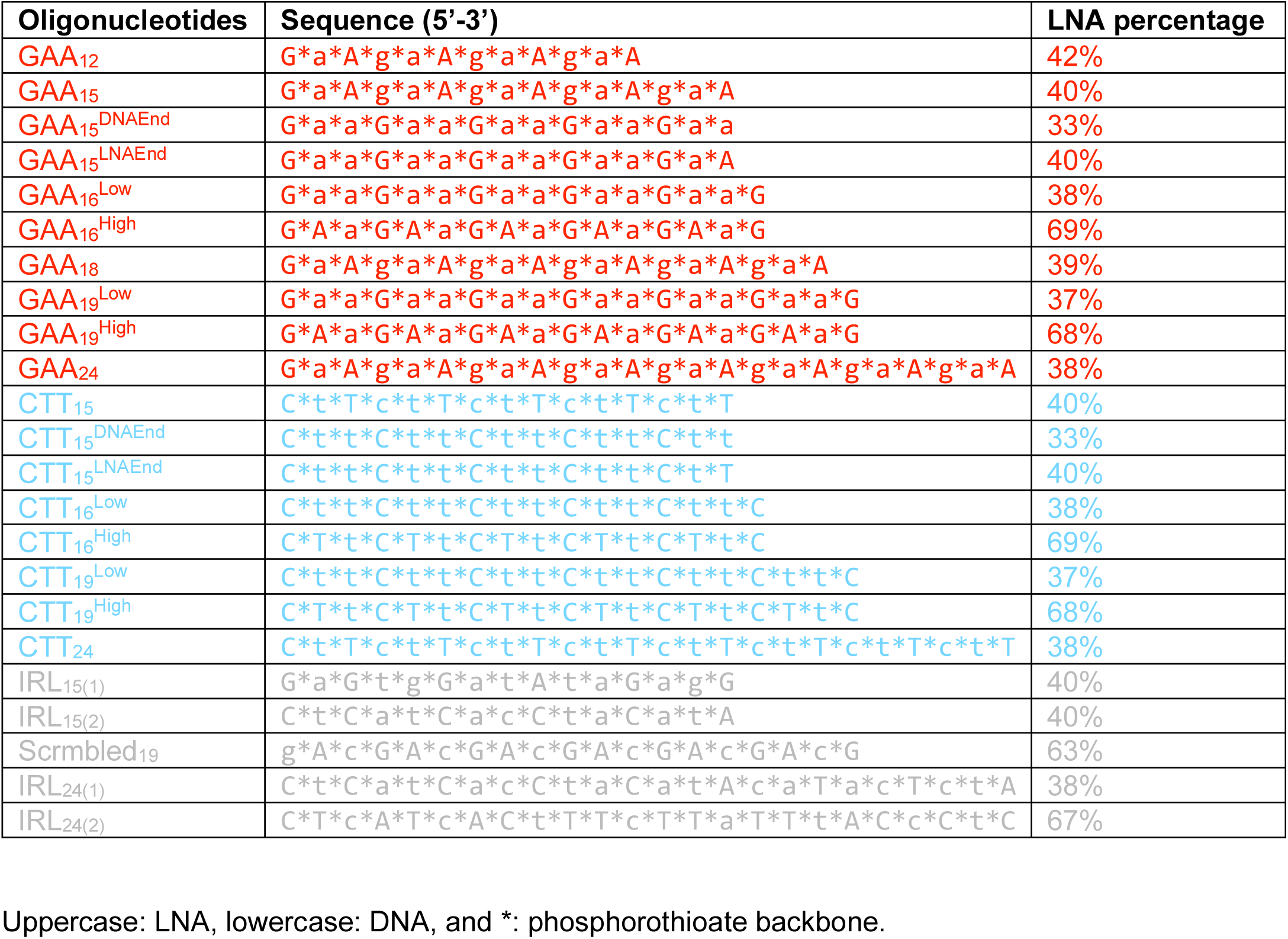
Oligonucleotides (ONs) used in this study.

**Figure 1.**
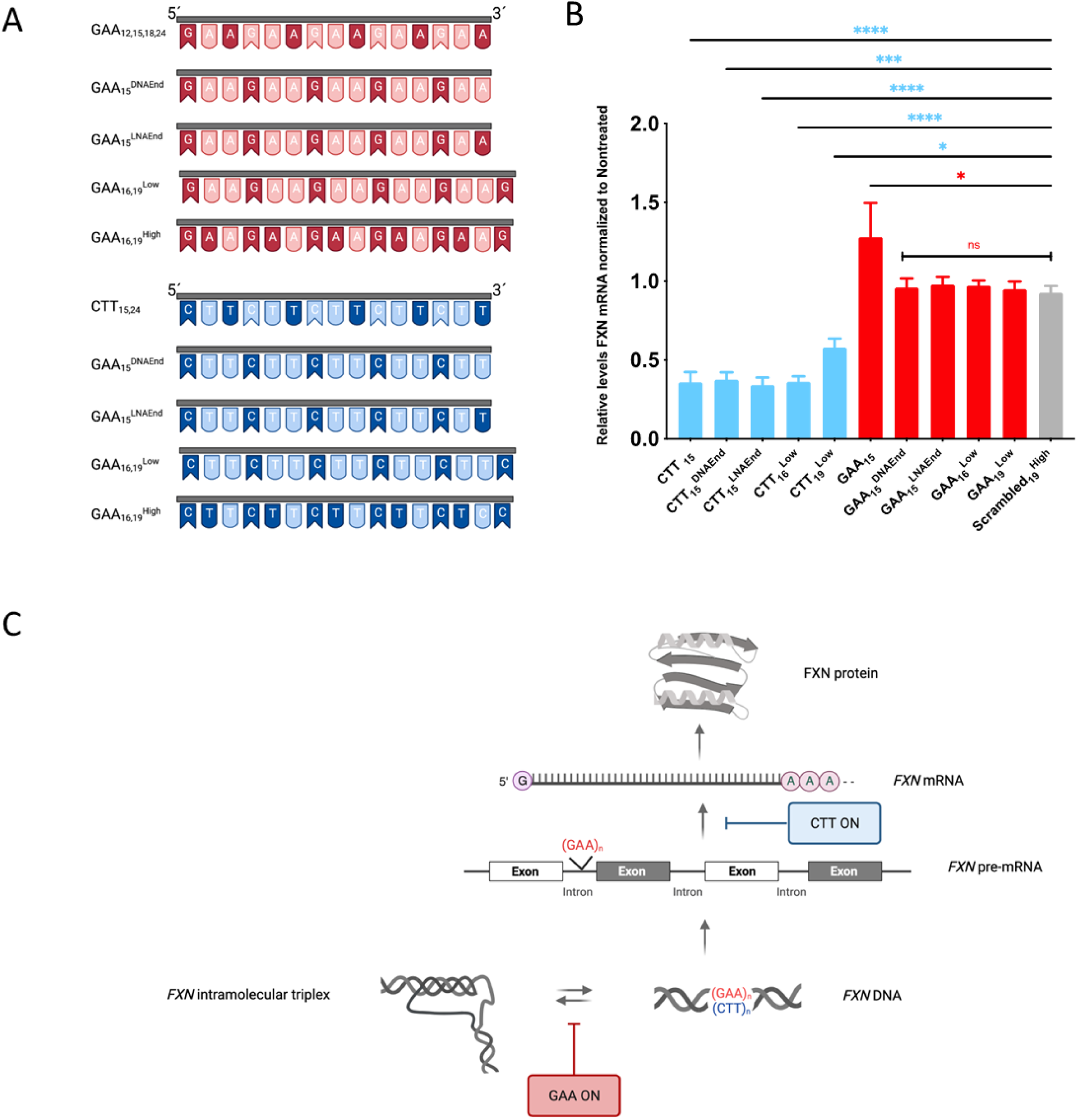
LNA composition influences the effect of GAA and CTT ONs on *FXN* mRNA expression. **A)** Illustrations of different GAA and CTT ON used in this study. LNA bases are in darker and DNA bases in lighter color. **B)** FA patient-derived fibroblasts carrying 330/380 GAA•TTC repeats (GM03816) were treated with 200 nM GAA AGOs and *FXN* mRNA levels were analyzed with RT-qPCR 4 days after transfection. Relative *FXN* expression of each treatment was compared to nontreated cells and normalized to the ratio of the *FXN* gene to *HPRT*. Results are presented as Mean with SEM (n=4). Statistical analysis was performed using one-way ANOVA, Multiple Comparison (Dunnett). (* = P < 0.05, ** = P < 0.01, *** = P < 0.001, **** = P < 0.0001; ns = nonsignificant).**C)** Schematic representation of GAA and CTT ONs interference with *FXN* expression.

Likewise, the corresponding CTT LNA/DNA mixmers were designed to be complementary to the released, single-strand chromosomal GAA region of the H-DNA or to the pre-mRNA (Table 1 and Figure 1A). To evaluate the activity of GAA and CTT ONs on *FXN* expression, the GM03816 patient-derived cells, carrying approximately 330/380 GAA•TTC repeats, were transfected with 200 nM of ONs and treated for 4 days. Data show that only GAA_15_ significantly upregulated *FXN* mRNA (Figure 1B). Additionally, the other two 15 mer GAAs (GAA_15DNAEnd_ and GAA_15LNAEnd_), both having DNA ‘adenosine’ instead of LNA ‘adenosine’ at the third position and either ending with a DNA or an LNA base, showed no significant effect on *FXN* mRNA expression (Figure 1B). Moreover, keeping the same design, LNA content and increasing the GAA ON length from 15 to 16 or 19 did not improve ON potency (GAA_16Low_^ON^; GAA ^Low^ Figure 1B). However, data shows that all the tested CTT ONs with the same features in terms of LNA substitution as the GAA AGOs, reduced *FXN* expression significantly (Figure 1B). We hypothesize that since GAA_15_ includes additional LNA modifications as compared to the other 15 mer GAAs tested, increasing the LNA content might enhance the efficacy of GAA AGOs to interfere with non-B-DNA targets. CTT ONs, on the other hand, could either hybridize to *FXN* pre-mRNA, thereby, sterically blocking the *FXN* expression; or bind directly to the single-strand GAA in the H-DNA formed at the GAA•TTC repeat in the *FXN* gene and therefore generate a more stable triplex structure on the opposite (template) strand preventing transcription elongation.

### Oligonucleotide length and LNA content determine the efficiency of GAA AGOs in upregulating ***FXN* transcription**

Subsequently, we tested if increasing the ON LNA content would affect *FXN* expression by promoting their GAA hybridizing capacity ^50^. Therefore, ONs were designed and synthesized to contain 68-69% LNA (Table 1). To study their activity, 200 nM of GAA and the corresponding control sequence, IRL_24(2)_, were transfected into GM03816 cells. *FXN* gene expression was analyzed after 4 days. We observed that ONs with 68-69% LNA content were toxic at 200 nM, a concentration at which ONs with lower LNA content were not (data not shown). Moreover, an increase in CTT or GAA ON length and LNA content did not improve their biological efficiency (data not shown). Thus, the toxicity of GAA AGOs with high LNA content makes them unsuitable as therapeutic candidates in FRDA.

We proceeded by assessing the influence of GAA ON length on *FXN* upregulation. Variants of GAA_15_, GAA_18_ and GAA_24_ were designed and synthesized, maintaining the LNA content within 38-40% (Table 1). For comparison, two different lengths of CTT ONs (CTT_15_ and CTT_24_) with the same design features were also studied (Table 1). Patient-derived fibroblasts GM03816 were transfected with 200 nM of GAA, or CTT ONs or their corresponding irrelevant counterparts (IRL_15(1)_ and IRL_15(2)_; all with 38-40% LNA content). FXN mRNA levels were evaluated 4 days post-transfection, showing that an increase in length of GAA AGOs significantly enhanced their efficacy with respect to FXN activation, with GAA24 promoting a 1.8-fold increase of FXN mRNA (Figure 2). In contrast, increasing the CTT ON length did not significantly affect FXN downregulation capacity (Figure 2). These data confirm the effectiveness of GAA AGOs with 38-40% LNA content in *FXN* upregulation.

**Figure 2.**
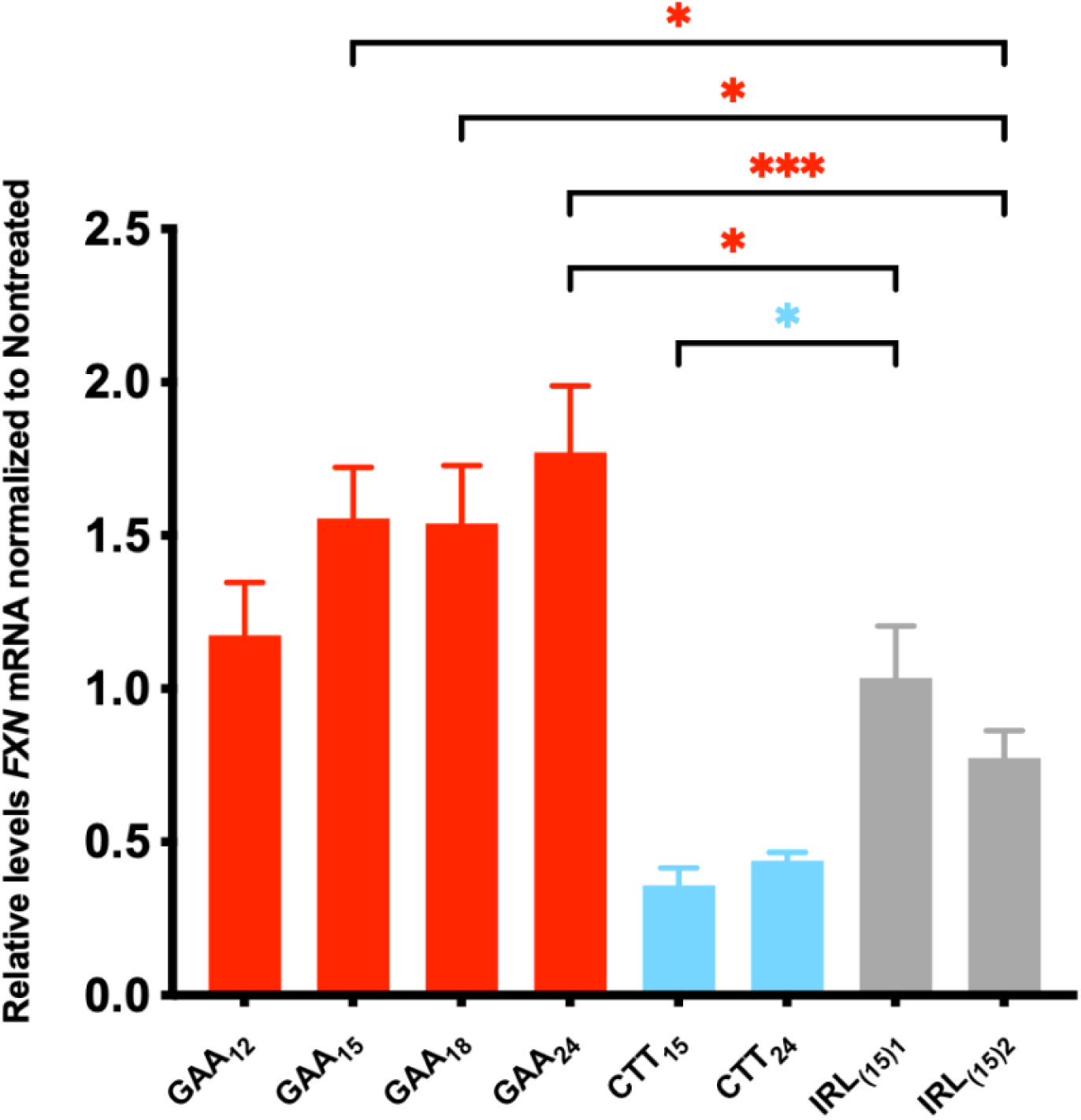
Increasing GAA ON length significantly improves *FXN* upregulation. FRDA patient-derived fibroblasts carrying 330/380 GAA•TTC repeats (GM03816) were treated with 200 nM GAA AGOs and *FXN* mRNA levels were analyzed with RT-qPCR 4 days after transfection. Relative *FXN* expression of each treatment was compared to nontreated cells and normalized to the ratio of the *FXN* gene to *HPRT*. Results are presented as Mean with SEM (n=3). Statistics were performed with one-way ANOVA toward IRL_15(1)_ (black *s) or to each other (blue *s). (* = P < 0.05, ** = P < 0.01, *** = P < 0.001, **** = P < 0.0001; no asterisk = not statistically significant).

Furthermore, the potency of GAA and CTT ONs in regulating *FXN* expression was also evaluated by gymnotic (naked) delivery in the presence of 9 mM CaCl_2_. This procedure has been found to yield a good correlation between *in vitro* and *in vivo* ON’s activities ^51^. For this purpose, ONs in a final concentration of 3 μM were added to the cells in a medium supplemented with 9 mM CaCl_2_. For comparisons, cells treated only with medium supplemented with 9 mM CaCl_2_, and nontreated cells (without ONs or CaCl_2_) were included as controls. Cells were harvested 4 days post-treatment, and *FXN* mRNA levels were determined using RT-qPCR. Contrary to expectations, the shorter GAA AGOs and GAA_15_ was significantly more potent for *FXN* mRNA upregulation in comparison to GAA_24_ at this particular concentration. Moreover, while both CTT_15_ and CTT_24_ downregulated *FXN* mRNA, we observed no statistically significant difference between them (Supplementary Figure 1).

### Longer GAA AGOs are more effective at lower doses

Due to the unanticipated trend where longer GAA AGOs showed a reduced effect on *FXN* expression in the gymnotic experiment as compared to transfection conditions and the corresponding shorter GAA AGOs, we decided to perform a dose-response of ONs when delivered naked in the presence of CaCl_2_^51^. ONs at final concentrations varying 32-fold, ranging from 0.1875 to 6 μM, were added to the medium supplemented with 9 mM CaCl_2_. For comparison, cells treated with 9 mM of CaCl_2_, and nontreated cells were included. Cells were harvested 4 days post-treatments, and *FXN* mRNA levels were determined using RT-qPCR. Interestingly, the GAA AGOs of varying lengths showed different dose-response curves. For GAA_15_, increasing the ON concentration to 6 μM translated to a more potent effect on *FXN* mRNA upregulation (Figure 3A). For GAA_18_, the effect was modest and seemed to plateau at concentrations above 0.75 μM (Figure 3B). In contrast, we observed that GAA24 was significantly more active at lower concentrations and displayed a maximum effect of 1.8-fold at 0.375 μM (Figure 3C). These results demonstrate the requirement of lower dosages for longer GAA AGOs for optimal *FXN* mRNA upregulation.

**Figure 3.**
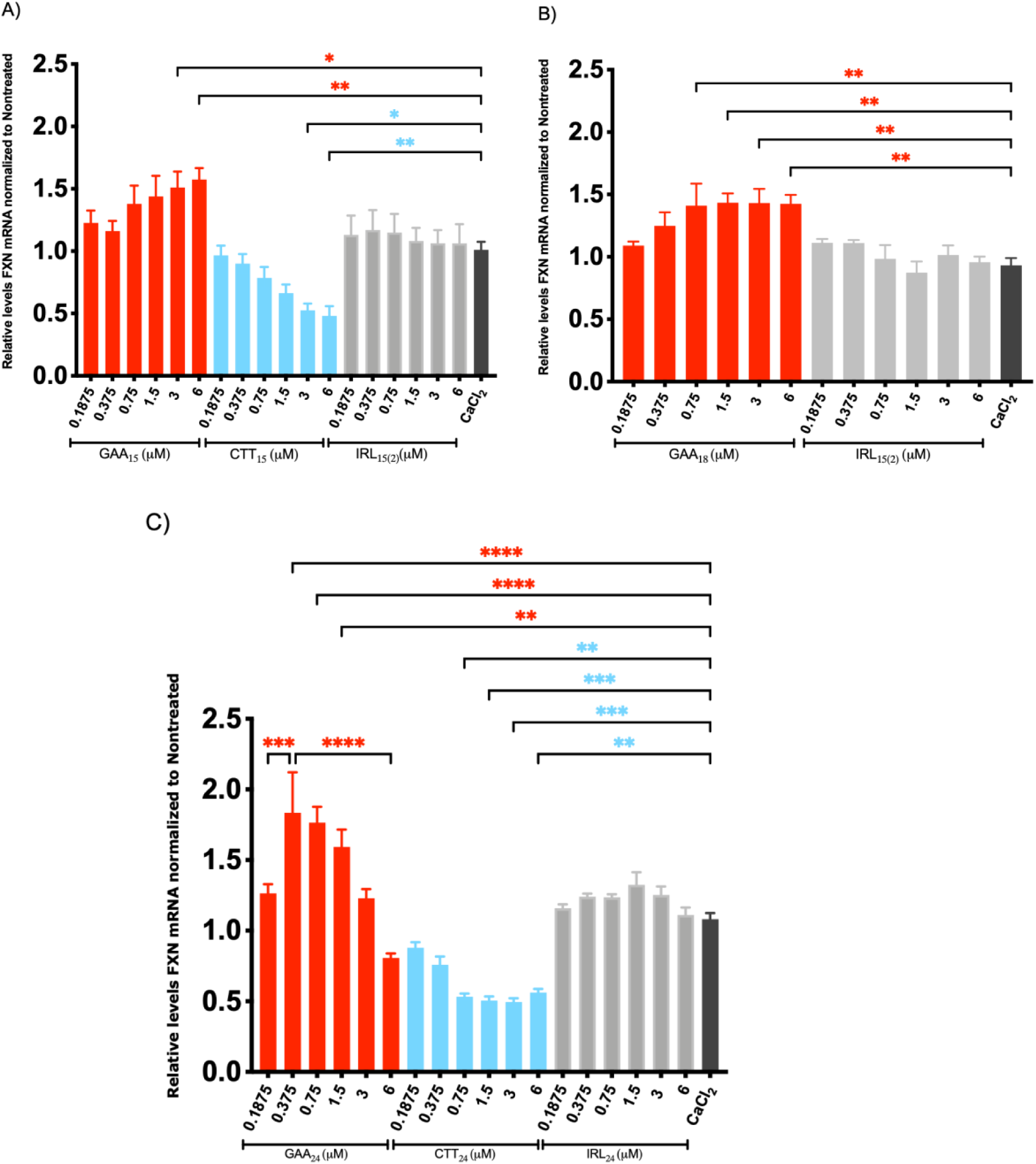
Dose-dependent *FXN* mRNA expression after gymnotic delivery of ONs. GM03816 cells were treated with ONs at concentrations ranging from 0,1875 to 6 µM in medium supplemented with 9 mM CaCl_2_. 4 days post treatments cells were harvested, and *FXN* mRNA levels were analyzed. The values were normalized to *HPRT* as reference gene and compared to nontreated cells without the presence of CaCl_2_. Results are presented as Mean with SEM (n ≥3). Statistics were performed with two-way ANOVA Multiple Comparison (Dunnett) towards CaCl_2_ only treated cells. (* = P < 0.05, ** = P < 0.01, *** = P < 0.001, **** = P < 0.0001 and ns = nonsignificant). **A)** GAA_15_ is more potent at *FXN* upregulation at higher doses, when comparing GAA_15_ at 6 versus 0.375 µM a significant difference is found. Similarly, CTT_15_ is significantly more effective at downregulate *FXN* mRNA at higher (6 µM) compared to lower (0,1875 µM) concentrations. **B)** GAA_18_ is significantly more effective In *FXN* mRNA upregulation at higher (0.75,1.5, 3 and 6 µM) compared to lower (0.1875 µM) doses. No significant difference between 0.75,1.5, 3 and 6 µM concentrations were found. **C)** GAA_24_ is significantly more potent in *FXN* mRNA upregulation at lower concentration (0,375 µM) compared to higher (6 µM). In contrast, CTT_24_ is significantly more active at higher (6 µM) versus lower (0.1875 µM) concentrations.

On the other hand, no significant upregulation of *FXN* mRNA was detected when treating FRDA patient-derived cells with GAA_16_^High^ and GAA_19_^High^ (both with 69% LNA content; Figure 4A and 4B).

**Figure 4.**
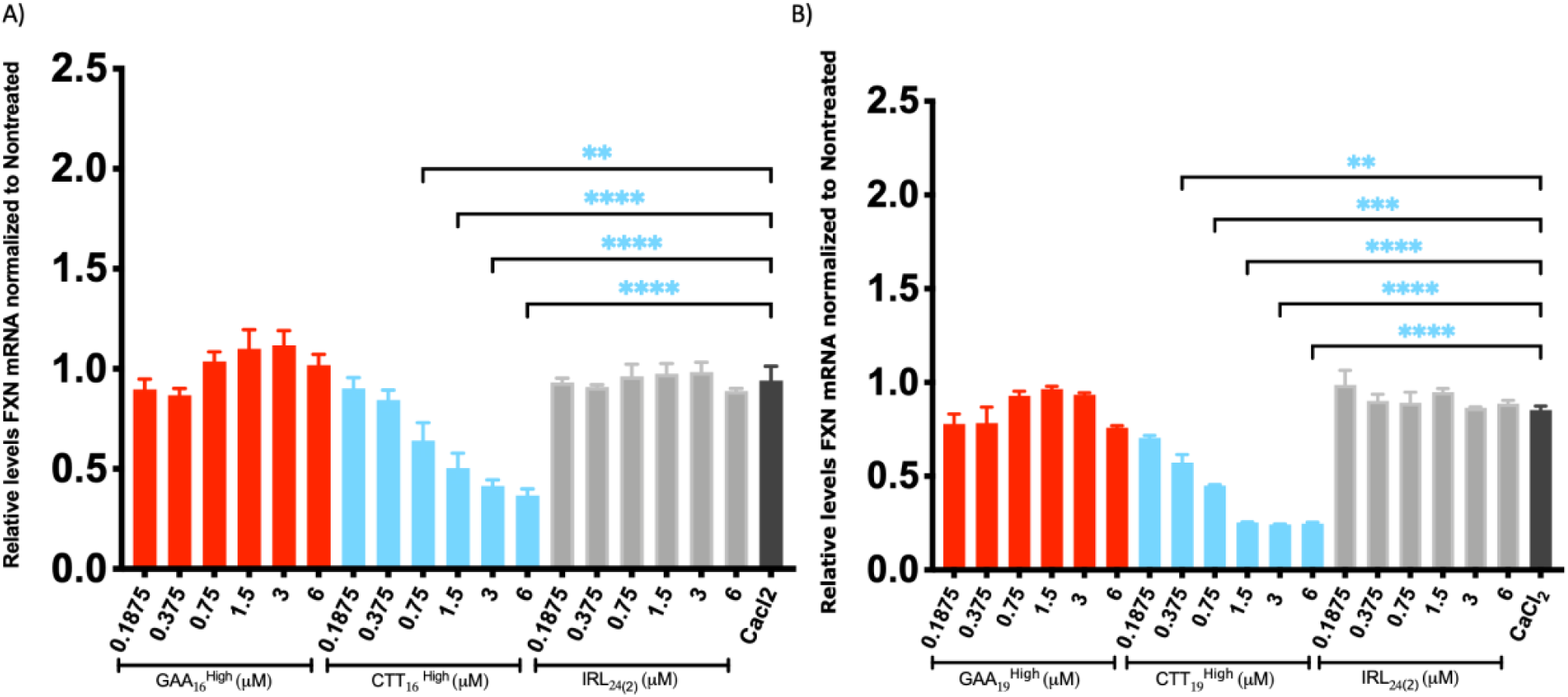
Dose response study of *FXN* mRNA expression after gymnotic delivery of ONs. GM03816 cells were treated with ONs at concentrations ranging from 0,1875 to 6 µM in medium supplemented with 9 mM CaCl_2_. 4 days post treatments cells were harvested and *FXN* mRNA levels were analyzed. The values were normalized to *HPRT* levels as reference gene and were compared to nontreated cells. Results are presented as Mean with SEM. Statistics were performed with one-way ANOVA Multiple Comparison (Dunnett), toward CaCl_2_ only treated cells. (* = P < 0.05, ** = P < 0.01, *** = P < 0.001, **** = P < 0.0001; no asterisk = not statistically significant). Both GAA ^High^ **(A)** and GAA ^High^ **(B)**, showed no significant effect compared to control on *FXN* upregulation in all tested concentrations. Whereas in CTT^High^ **(A)** and CTT ^High^ **(B)** a dose dependent and significantly downregulation of *FXN* mRNA at 6 µM compared to 0,1875 µM was found.

The CTT ONs were used as control, since it was established in the transfection experiments that these resulted in the opposite effect, thereby, reducing *FXN* mRNA levels. The silencing effect of CTT_15_,CTT _High, CTT_ ^High^ and CTT_24_ showed a different dose-response for the concentration interval 0.1875 to 6 μM, with significantly greater *FXN* downregulation at higher concentrations (Figures 3A, 3B, 4A and 4B). Furthermore, selected ONs with 40% LNA content were shown to be nontoxic as determined by the WST-1 viability assay at all tested concentrations (Supplementary Figure 2). Based on these results, increasing the LNA content of GAA AGOs reduces their activity above a certain threshold of 3 to 6 µM in a gymnotic context. Moreover, the longer the GAA ONs, the more effective they were at lower doses. Our findings imply the potential benefits of ONs in the treatment of FRDA and also suggest that there is an intricate balance between dose and length with regard to the therapeutic activity of AGOs.

### GAA AGOs upregulate FXN protein expression

Following the evaluation of the optimal length and sequence of GAA AGOs in *FXN* mRNA upregulation by RT-qPCR, we aimed to assess if that effect can be translated to protein production. FRDA fibroblast cells GM03816 were transfected with 100 nM of the ONs with 40% LNA content (GAA_15_, GAA_18_, GAA_24_, CTT_15_, CTT_24_, IRL_15(1)_, IRL_15(2)_ and IRL_24(1)_). The cells were harvested after 4 days and protein expression was analyzed by western blotting. Similar to mRNA experiments, both CTT_15_ and CTT_24_, significantly downregulated FXN protein expression compared to cells treated with IRL ONs (Figure 5 A and B). However, GAA_24_ treatment showed a significant increase in FXN protein levels which relates to the increased levels of *FXN* mRNA. Because 24-nucleotide long GAA AGOs resulted in a superior effect on FXN protein upregulation, GAA_24_ and the corresponding control ONs, IRL_15(2)_, and CTT_24_ were gymnotically delivered to GM03816 cells, in the presence of 9 mM CaCl_2_, at concentrations ranging from 0.375 to 1.5 μM. In agreement with our previous findings regarding the effect on *FXN* mRNA levels, our data confirmed that the GAA_24_ AGO significantly upregulates FXN protein expression under gymnotic ON delivery conditions. Moreover, cell treatment with the CTT_24_ control ON reduced FXN protein levels following the previous trend where increased concentration was more efficient (0.375 μM and 1.5 μM, respectively), in line with the results obtained by analysis of mRNA expression levels (Figure 5 C and D).

**Figure 5.**
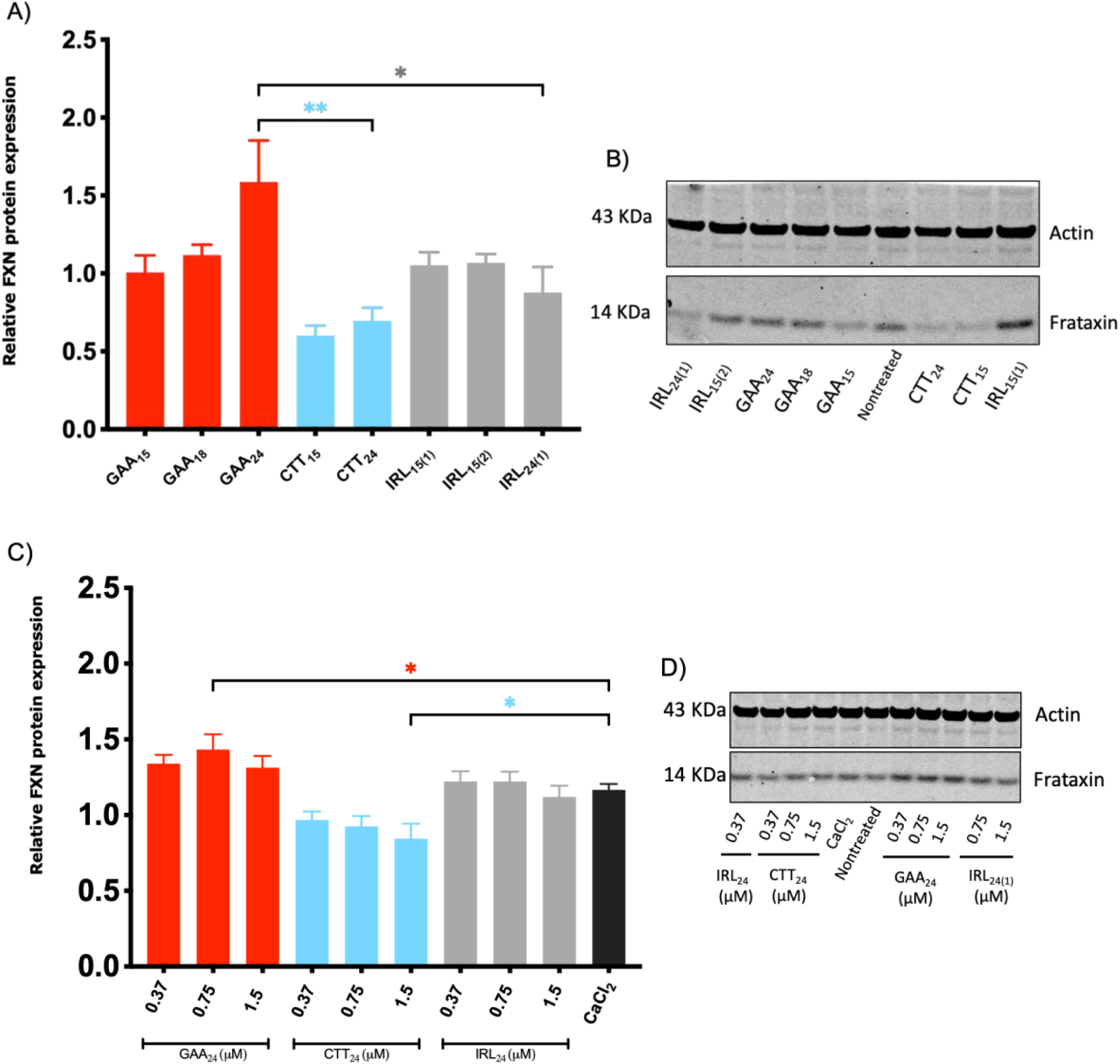
Effect of GAA AGOs on FXN protein upregulation. **A)** FXN protein levels were measured with western blotting 4 days after the gymnotic delivery of GAA_24_, CTT_24_ and IRL_15(2)_ in medium supplemented with 9 mM CaCl_2_. The values were normalized to Actin levels as reference gene and compared to nontreated cells. Results are presented as Mean with SEM (n =3). Statistics were performed with two-way ANOVA Multiple Comparison, (Dunnett), toward CaCl_2_ only treated cells (* = P < 0.05, ** = P < 0.01, *** = P < 0.001, **** = P < 0.0001; no asterisk = not statistically significant). **B)** Representative western blot gel of gymnotic delivery of GAA_24_, CTT_24_ and IRL_15(2)_ in medium supplemented with 9 mM CaCl_2_, 4 days after treatments. **C)** Effect of ONs with 40% on FXN protein expression after 4 days transfection of 100 nM ONs with Lipofectamine LTX. The values were normalized to Actin levels as reference gene and were compared to nontreated cells. Results are presented as Mean with SEM (n ≥3). Statistics were performed with one-way ANOVA Multiple Comparison (Dunnett) (* = P < 0.05, ** = P < 0.01, *** = P < 0.001, **** = P < 0.000; no asterisk = not statistically significant). **D)** Representative western blot gel of transfection of ONs with 40% LNA 4 days after treatment.

### GAA AGOs upregulate *FXN* mRNA expression in additional FRDA cell model with a higher repeat number

The majority of FRDA patients carry 600 – 900 repeats in *FXN* gene ^7^, with a maximum reported to this date of 1700 repeats ^3^. After confirming GAA AGOś efficiency in enhancing *FXN* mRNA expression in an FRDA cell model carrying 330/380 GAA•TTC repeats, we hypothesized that AGOs could act similarly in a more clinically relevant cell model of the disease. FRDA patient-derived fibroblasts GM03665 carrying approximately 780/1410 GAA•TTC repeats were transfected with 200 nM of GAA and CTT ONs. To assess how the length affects ONs potency, different lengths of GAA AGOs were tested, namely 12-, 15-, 18- and 24-mers. Selected cognates CTT_15_ and CTT_24_ and AGO’s corresponding controls IRL_15_ and IRL_24_ were also used. Regarding LNA content, in the lower number of repeats GM03618 fibroblasts the best-performing ON design included 38-40% LNA. Based on this, GM03665 fibroblasts were treated with different ONs containing 38-40% LNA content. As a control, nontreated cells were also included. Cells were harvested 4 days after treatments, and *FXN* mRNA levels were assessed by RT-qPCR (Figure 6). A similar experiment was performed at a concentration of 100 nM (Supplementary Figure 3).

**Figure 6.**
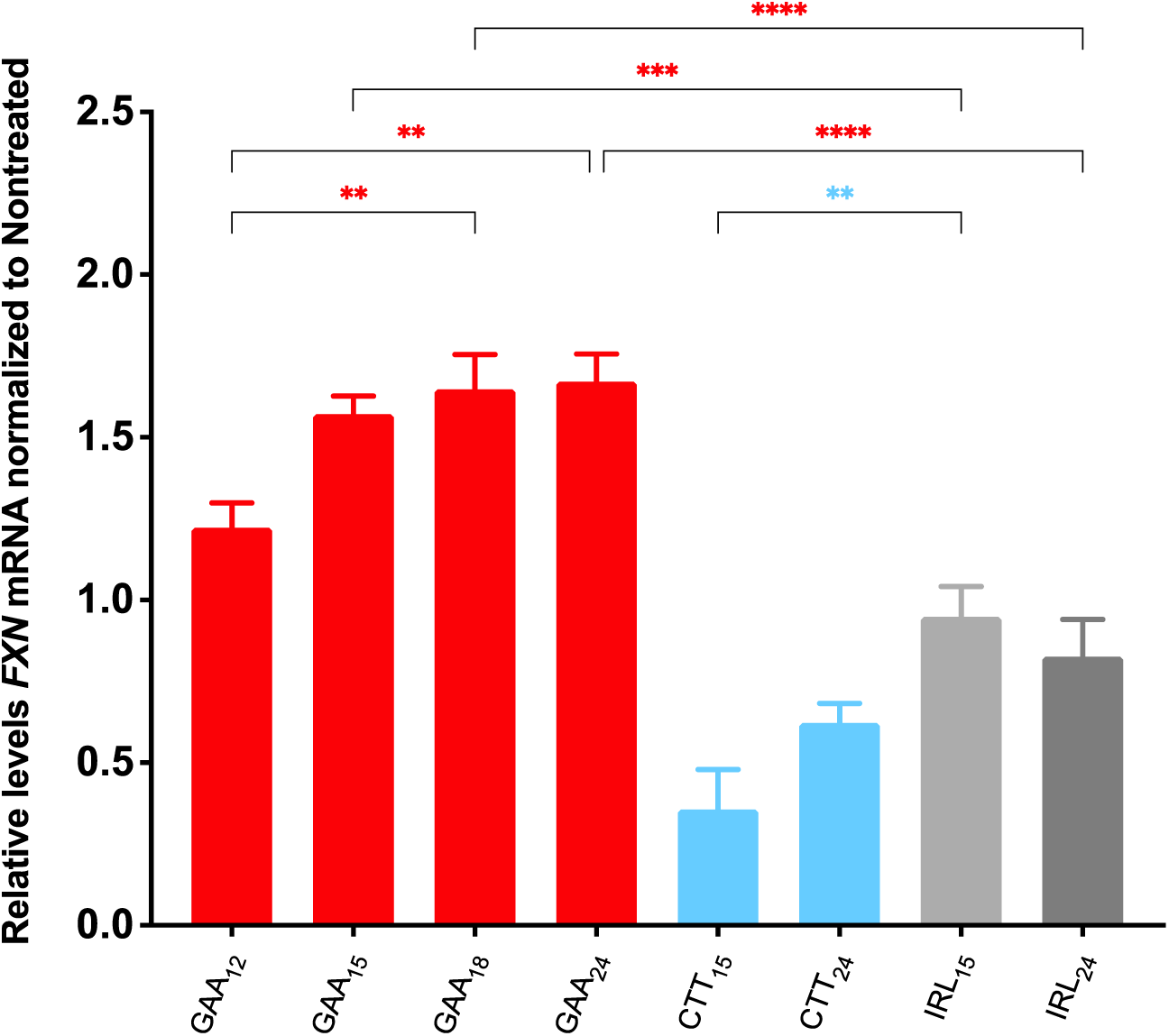
Increasing GAA AGOs length enhances *FXN* upregulation in a FRDA cell model with a higher number of repeats. GM03665 fibroblasts were treated with ONs of different lengths of GAA, CTT and IRL at 200nM. Treated and nontreated cells were harvested 4 days after transfection, and *FXN* mRNA levels were assessed by RT-qPCR. The values were normalized to *HPRT* levels as reference gene and were compared to nontreated cells. Results are presented as Mean with SEM (n ≥3). Statistics were performed with one-way ANOVA Multiple Comparison, (Dunnett), towards nontreated cells. (* = P < 0.05, ** = P < 0.01, *** = P < 0.001, **** = P < 0.0001; no asterisk = not statistically significant). GAA_15,_ GAA_18_ and GAA_24_ significantly upregulate *FXN* mRNA compared to nontreated cells. CTT_15_ resulted in significant downregulation of *FXN* mRNA.

The results show that *FXN* mRNA levels are significantly upregulated when GM03665 fibroblasts are transfected with GAA_15_ and GAA_24_ at 200 nM (Figure 6). Similarly to what was observed in GM03816 fibroblasts, the obtained *FXN* upregulation reached a maximum effect of 1.6-1.7 fold increase. Again, the enhanced upregulation correlated positively with the ON length, as increasing GAA ON length significantly potentiated a higher fold of *FXN* upregulation. The cognates CTT_15_ and CTT_24_ ON controls resulted, as expected, in significant downregulation of *FXN* mRNA. These data validate the effectiveness of AGOs in the context of different FRDA patient-derived fibroblasts containing a higher number of expanded GAA•TTC repeats.

### Gymnotic delivery of GAA_24_ AGOs successfully upregulates *FXN* mRNA expression in the higher-repeats cell line

In the lower number of repeats GM03618 fibroblasts, the best-performing combination regarding ON length and LNA content, was the 24 mer GAA ON containing 38-40% LNA content. Similarly, GM03665 fibroblasts transfected with GAA_24_ containing the same LNA amount resulted in the highest *FXN* upregulation. Based on this, GM03665 fibroblasts were treated with GAA_24_ containing 38-40% LNA in the medium enriched with 9 mM CaCl_2_, at final concentrations ranging from 0.18 to 3 μM. A similar treatment was carried out using the corresponding CTT_24_ control ON. For the irrelevant ON, IRL_24_, only selected concentrations were used. Nontreated cells were also included. Cells were harvested 4 days after treatments, and *FXN* mRNA levels were assessed by RT-qPCR.

Cell treatment with GAA_24_ showed a dose-response and the AGO significantly upregulated *FXN* mRNA levels at 0.75 μM and 1.5 μM, with a maximum effect of a 1.5-fold increase (Figure 7). This pattern is similar to what was observed in GM03816 fibroblasts where lower concentrations of GAA_24_ led to optimal *FXN* mRNA upregulation. The cognate CTT_24_ induced significant downregulation of *FXN* mRNA levels for 0.18, 0.75 and 1.5 μM, in a dose-response manner (Figure 7). Hence, CTT_24_ behaved here again in an opposite manner compared to GAA_24_, being significantly more potent at higher concentrations (3 µM) compared to lower concentrations (0.18 µM)., which is in accordance with our findings observed in the GM03816 fibroblasts. Altogether, these data confirm the efficiency of gymnotically delivered GAA AGOs in *FXN* upregulation in the context of a more severe cell model of FRDA.

**Figure 7.**
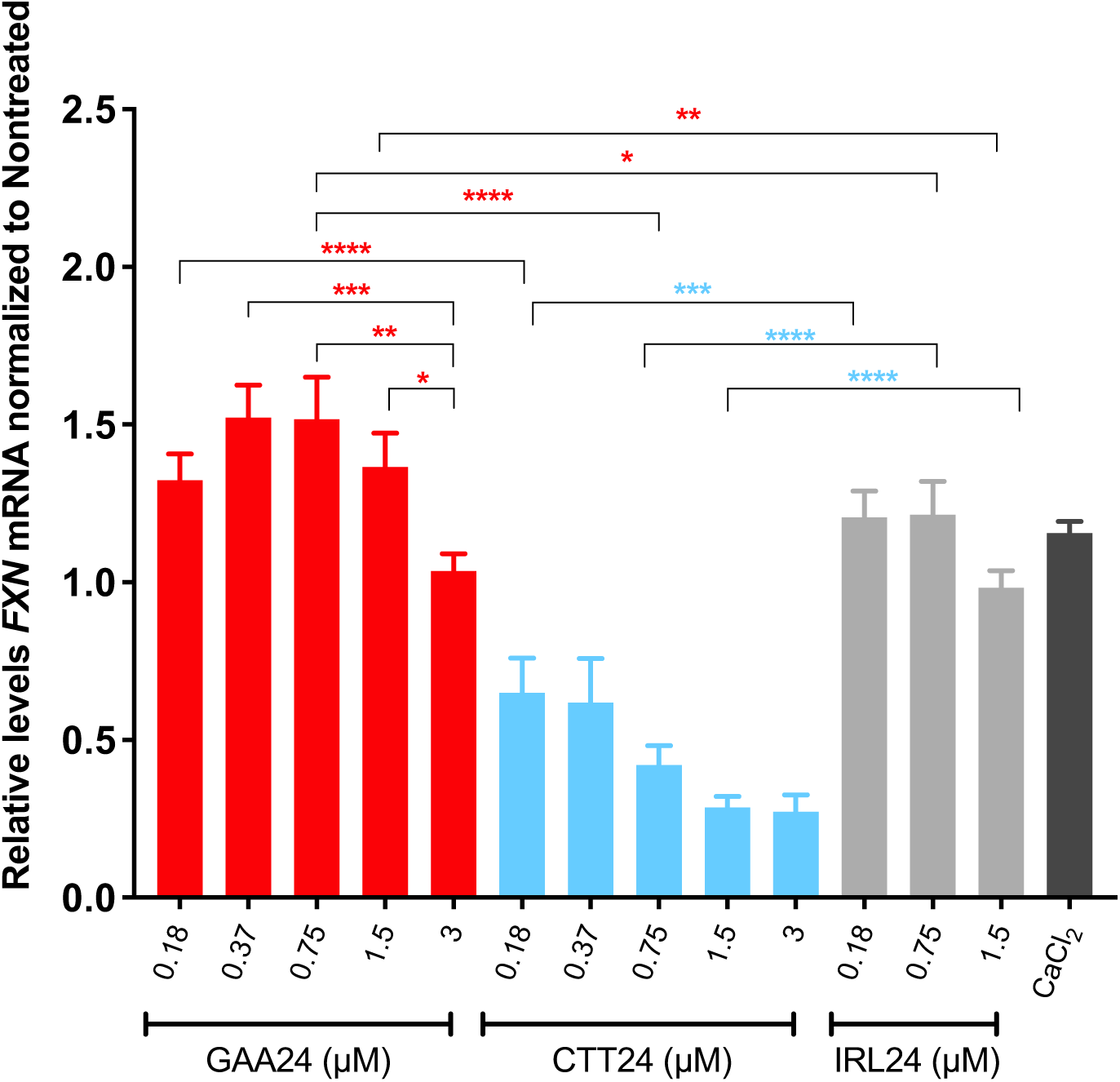
Dose response study of *FXN* mRNA response after gymnotic delivery of GAA_24_ ONs. GM03665 fibroblasts were treated with of GAA_24_ and CTT_24_ ONs at concentrations ranging from 0.18 to 3 µM in medium supplemented with 9 mM CaCl_2_. For IRL_24_, only selected concentrations were used. Treated and nontreated cells were harvested 4 days post-treatments, and *FXN* mRNA levels were assessed by RT-qPCR. The values were normalized to *HPRT* levels as reference gene and were compared to nontreated cells. Results are presented as Mean with SEM (n ≥3). Statistics were performed with two-way ANOVA Multiple Comparison (Tukey). (* = P < 0.05, ** = P < 0.01, *** = P < 0.001, **** = P < 0.0001; no asterisk = not statistically significant). Both 0.75 and 1.5 µM of GAA_24_ showed significant effect compared to their respective IRL_24_ controls on *FXN* upregulation. 0.18, 0.75 and 1.5 µM of CTT_24_ resulted in significant downregulation *FXN* mRNA compared to their respective IRL_24_ controls.

## DISCUSSION

In this study we designed modified, single-strand AGOs to target a non-B DNA structure formed at *FXN*-expanded GAA•TTC repeats as a potential therapeutic approach for FRDA. We designed several PS modified DNA/LNA mixmers with varying length, LNA content and LNA pattern. We assessed their efficiency in two FRDA-patient derived cell models containing different GAA•TTC repeats number.

Using both lipid-based transfection and gymnosis, treatment with the GAA AGOs led to a significant increase in FXN mRNA and protein levels. However, this was only observed when adding specific number of LNA modifications in certain positions within the sequence, showing that the design of the GAA AGO is crucial for its efficacy. Contrary to the GAA AGOs upregulating effect, CTT ONs consistently showed significant downregulation of *FXN* mRNA.

In FRDA the expanded GAA•TTC repeats at the *FXN* locus leads to the formation of non-canonical DNA (H-DNA) rather than the common, intrinsic B-DNA structure. The expanded GAA•TTC repeats at intron 1 of the *FXN* gene, form a parallel ^46^ or antiparallel triplex ^52^. The parallel triplex is formed by the disruption of one part of the double helix containing (GAA)_n_•(TTC)_n_ expanded repeats, and, while the CTT strand of the duplex folds back and makes hydrogen bonds with the undisrupted part of the duplex, the GAA strand remains a single-strand ^53^. The formation of an intracellular triplex leads to RNA polymerase blockage and eventually transcriptional silencing of *FXN* mRNA expression ^54^.

As the pathogenic, expanded GAA•TTC repeats are located in an intronic region of the *FXN* gene, the use of sequence-specific AGOs targeting chromosomal DNA has great potential. In contrast to the conventional approach of ASOs, AGOs are designed to specifically target and modulate gene expression at the chromosomal level. Unlike RNA targets, which are constantly produced, there are only two chromosomal regions in the *FXN* gene to which the AGOs can bind, which would be beneficial since lower doses are needed in this case.

The GAA AGOs are expected to bind and disrupt the formation of a triplex/H-DNA structure leading to enhanced transcription elongation and upregulation of *FXN* gene expression. This has been validated previously in a study that we carried out using chemical and structural probing of the DNA complexes formed in the presence of modified ONs ^45^. Cognate CTT ONs with the same design and length as the GAA AGOs were also explored. CTT ONs are believed to bind sequence-specifically to the formed single strand (GAA)_n_ at the FXN gene, making the H-DNA more stable. Additionally, they can sterically block the pre-mRNA ^54^. Furthermore, we recently showed that both GAA and CTT ONs reduce GAA•TTC expansion frequency in an experimental model system ^47^. To follow up on these two studies, we further optimized these ONs and assessed their potency in FXN upregulation, a key step in the development of FRDA therapeutics.

Targeting and disrupting an intracellular triplex is complex, especially in a region which also is believed to be epigenetically inactivated ^54^. We therefore evaluated the effect of GAA AGOs on *FXN* levels by varying their lengths and LNA content. Increasing the LNA proportion had no beneficial effect on *FXN* mRNA upregulation, which could be explained by the fact that there is a need for a balance between GAA ON invasion with subsequent binding and disruption of the triplex, followed by AGO disassociation from DNA, thereby enabling the RNA polymerase to elongate. This suggests that since LNA-modified ONs show greater affinity and stability ^55^, their sequence composition also needs to be well-adjusted. We found that increasing the length of GAA AGOs with an optimized LNA content enhanced *FXN* mRNA upregulation in a dose-dependent fashion. Thus, GAA_24_ holds better promise for *FXN* upregulation at lower concentrations, rendering it of interest as a potential therapeutic agent for the treatment of FRDA. Having examined this in two cell lines originating from patients carrying different full-penetrance allele sizes, we observed that the maximum upregulation is reached at the same concentration range in both cell lines. The levels of upregulation we achieved is physiologically relevant as in FRDA the heterozygous carriers express 50% less FXN while developing an atypical form of FRDA with almost no phenotype and low disease progression ^19,56^. Furthermore, the level of FXN upregulation needs to be well adjusted as it has been shown that overexpression of FXN causes toxicity ^57^. These results could indicate the same optimal dosage for FRDA patients independently of their allele sizes. Furthermore, a significant *FXN* mRNA upregulation was achieved by shorter ONs (e.g. GAA_15_), which is also of interest in a treatment context, since *in vivo* uptake of AGOs may be length-dependent. The results using GAA AGOs are therefore in agreement with our hypothesis that the balance between the capacity of GAA AGOs for invasion/disruption of the triplex and subsequent disassociation from genomic DNA needs careful optimization with respect to design and dosage.

Moreover, in contrast to a mechanism whereby the H-DNA is dissolved by GAA AGOs, for CTT ONs an effect on DNA as well as on pre-mRNA is likely. Irrespective of the mechanism, CTT ONs with enhanced hybridization properties, hence higher LNA content, significantly reduced the *FXN* mRNA expression regardless of LNA content and its position. This was shown in both FRDA patient cell lines.

A recent finding further illustrates that the enhanced expression from the *FXN* gene upon ON treatment differs from the mechanism underlying the corresponding pathognomonic repeat expansions in FRDA. Hence, while GAA-, but not CTT ONs enhance *FXN* gene expression in FRDA patient-derived cell lines, GAA as well as CTT ONs block the expansion of GAA•TTC repeats in reporter cells ^47^.

FRDA is not the only example of experimental AGO treatment in repeat expansion disorders. Thus, we have recently reported that CAG•CTG repeat-targeting AGOs had a distinct effect on the *HTT* locus, whose repeats are expanded in Huntington’s Disease (HD). *HTT* mRNA and protein were *not* upregulated, but instead significantly reduced in HD patient-derived fibroblasts and neural stem cells differentiated from induced pluripotent stem cells ^58,59^. Since the underlying mechanism of experimental AGO therapy is expected to differ between FRDA and HD, there are some important variables to consider. For the *HTT* locus, a high LNA content was essential for obtaining an effect on expression since AGOs carrying 31.5% LNA were inefficient ^60^. Unlike our findings in FRDA, AGOs containing ca 60% LNA were highly efficient in HD patient-derived cell lines. Importantly, the non-B DNA structure of the affected genes is not the same in FRDA and HD. In the *HTT* locus, the conformation is considered to exist as a hairpin, whereas in the *FXN* gene, a triplex/H-DNA is formed ^61^.

While we examined several different ON constructs, compared to the extensive testing of compounds generally performed in the pharmaceutical industry, our ON catalogue has been modest in size. Continued optimization of these AGOs may therefore yield even more efficient lead compounds to be developed as therapeutics. Nevertheless, we demonstrated that the AGO targeting concept is valid in two FRDA patient-derived cell lines carrying different GAA•TTC repeat expanded alleles. The AGOs that have been previously reported to disrupt H-DNA formation, now show for the first time the capacity to significantly upregulate *FXN* expression. This suggests that they can potentially be used as a treatment option for FRDA.

## METHODS

### Oligonucleotides

LNA/DNA mixmers were purchased from Eurogentec S.A. or were synthesized at the Nucleic Acid Center, University of Sothern Denmark. The ONs were purified by reversed-phase HPLC, and their composition confirmed by MALDI-TOF mass spectrometry. The GAA AGOs were designed to target the pyrimidine motif triplex formed at the *FXN* intron expanded GAA•TTC repeats while the CTT ONs were designed to bind to the single-stranded GAA region of the repeats. Control ONs were designed with the same LNA composition and percentage. The complete list of AGOs used in this study is presented in Table 1.

### Cell culture, transfection and gymnosis

The primary fibroblasts derived from FRDA patients were obtained from the Coriell Institute. The GM03816 cells contain approximately 330/380 GAA•TTC repeats and the GM03665 cells contain approximately 780/1410 GAA•TTC repeats. Fibroblasts were maintained in a humidified incubator at 37 °C with 5% CO_2_. Cells were grown in DMEM (Dulbecco’s Modified Eagle Medium) with pyruvate and low glucose (Gibco), supplemented with 15% Fetal Bovine Serum (FBS) (Gibco). Lipofectamine LTX with PLUS reagent (Invitrogen by Thermo Fisher Scientific) was used to transfect the ONs according to the manufacturer’s recommended protocol. Briefly, cells were seeded a day before transfection at 1 x 10^4^, 8 x 10^4^, or 3 x 10^5^ cells per well in 96, 24 or 6-well plates, respectively. ONs were formulated with Lipofectamine LTX and PLUS reagent (Invitrogen by Thermo Fisher Scientific) at a final concentration of 100 or 200 nM in OptiMEM reduced serum medium (Gibco). For gymnosis experiments, ONs were added freshly to the medium supplemented with 9 mM CaCl_2_ the day after seeding according to the Hori et al. protocol ^51^. Cells were lysed for RNA isolation or protein expression evaluation 4 days later.

### RNA isolation and reverse-transcription quantitative PCR

Total RNA was isolated with Tri-reagent® (Sigma-Aldrich) or RNeasy plus kit (QIAGEN) according to the manufacturer’s protocol. Quantity and quality of RNA was measured with NanoPhotometer (Implen). 200 ng of total RNA was used for cDNA synthesis with the High-Capacity cDNA Reverse Transcription Kit using random primers (Applied Biosystems). RT-qPCR was performed by the CFX96 Real-Time PCR system (Bio-Rad) using TaqMan Fast Advanced Master Mix (Applied Biosystems) with 20 ng of cDNA as a template. *FXN* Exon4-Exon5 was amplified using the following primers and probe: FXN-sense: 5’-gtggagatctaggaacctatg, FXN-antisense: 5’-ttaaggctttagtgagctctg and FXN-probe: 5’-[HEX]tccagtcataacgcttaggtccac[BHQ1] ^62^. Normalization was performed using the following primers and probe for hypoxanthine phosphoribosyltransferase 1 (*HPRT1*) gene: HPRT-sense: 5’-agggatttgaatcatgtttg, HPRT-antisense: 5’cgatgtcaataggactcc, and HPRT-probe: 5’-[6FAM] actcaacttgaactctcatcttaggct[BHQ1]. The data were analyzed with CFX Maestro software (Bio-Rad) using the ΔΔCq method.

### Western blotting

Cells from 6-well plates were trypsinized (Gibco) and collected in Eppendorf tubes. Cells were lysed with RIPA buffer for 30 min on ice following centrifugation at top speed for 15 min at 4 °C. 1x Concentrated sample reducing agent (Invitrogen), and NuPAGE LDS sample buffer (Invitrogen) was added to the supernatant after centrifugation. The samples were heated at 75 °C for 10 min prior to loading on the gel. Proteins were separated on Bolt™ 4 to 12% Bis-Tris gels (Invitrogen) at voltage 70 for 20 min following 90 min at voltage 130. Gels were transferred onto nitrocellulose membranes (Invitrogen) using iBlot2 Gel Transfer Device (Invitrogen). The membranes were blocked with Odyssey Blocking Buffer (TBS) (LI-COR Biosciences) for 1 h. Blocked membranes were probed with anti-FXN primary antibody (ab110328, Abcam) (1:500) and anti-Actin (1:10^6^) (Sigma-Aldrich) as a reference. The primary antibodies were diluted in a 1:1 ratio of PBST (Phosphate-buffered saline with 0.1% Tween 20) and blocking buffer and incubated at 4 °C on a rocking plate shaker overnight. After primary antibody incubation, the membranes were washed 5 times for 5 min at room temperature with 1x PBST and then incubated with IRDye 800Cwgoat anti-mouse IgG (1:40,000) as a secondary antibody (LI-COR) for 1 h at room temperature. Membranes were washed 5 times for 5 min at room temperature with 1x PBST and 2 x 3 min with PBS prior to scanning. The signals were detected with an Odyssey imager (LI-COR) at 800 nm.

### Viability assay

To assess the viability of the cells upon ON treatments, the WST-1 assay (Merck) was used according to the manufacturer’s recommended protocol. Briefly, cells were cultured in a 96-wells plate at a final volume of 100 μL. The day after seeding, the cells were treated with the ONs as stated above. Two days after treatment, media was substituted with fresh media containing 10 μL (1:10 dilution) WST-1 reagent and incubated at 37°C with 5% CO_2_ for 2 h. The signals were measured with SpectraMax i3x (Molecular devices) at 450 nm with 600 nm as reference wavelength. The relative value of cell viability was calculated based on the ratio of treated and non-treated cells at 450 nm wavelength.

### Statistical analysis

Data are expressed as mean ± SEM. Statistical analyses were performed using One or Two-way ANOVA, Multiple Comparison (Dunnett) test using GraphPad Prism 6 Software (GraphPad, Inc). A probability of less than 0.05 was considered statistically significant.

## FUNDING and ACKNOWLEDGMENT

Funding was kindly provided from Hjärnfonden, The Swedish Research Council, Swelife-Vinnova, CIMED (Center for Innovative Medicine) and Region Stockholm (OS, NM, CIES, and RZ). This project has also received funding from the European Union’s Horizon 2020 research and innovation programme under grant agreement No 956070 (SM; granted to CIES). NovoNordisk [NNF21OC0072778] ’Pioneer Innovator 2-2021’ (NM, TU)

## AUTHOR CONTRIBUTIONS

All authors declare a contribution to this paper. RZ, CIES, and NM, designed and planned the study with input from PB, PTJ and JW. NM, SM, CV, FF and OS performed and analyzed experiments. JW and PTJ contributed to chemical synthesis. NM wrote the first draft of the manuscript. SM, CR, TU, PB, PJT, JW, CIES and RZ, took part in the revision of the manuscript for important intellectual content. All authors reviewed and approved the final version of the manuscript.

## DATA AVAILABILITY

The data that support the findings of this study are available from the corresponding author upon reasonable request.

## CONFLICT OF INTEREST STATEMENT

RZ has a granted patent for the treatment of Friedreich’s ataxia.

## Notes

### Competing Interest Statement

Rula Zain has a granted patent for the treatment of Friedreichs ataxia.

