## Supplementary material for "Anti-gene oligonucleotides targeting Friedreich’s ataxia expanded GAA•TTC repeats increase Frataxin expression"

**Supplementary information**

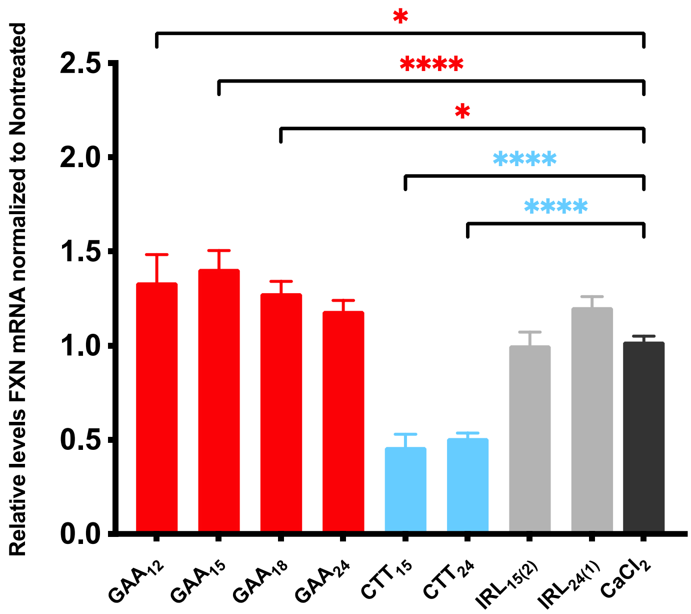


**Supplementary Figure 1.** 3 µM ONs in medium supplemented with 9 mM $CaCl_2$ were added to the cells the day after plating. After 4 days total RNA was extracted, mRNA was quantified with RT-qPCR, *FXN* levels were normalized to *HPRT* and compared to nontreated cells. Results are presented as Mean with SEM (n ≥3). Statistics were performed with one-way ANOVA Multiple Comparison (Dunnett) toward $CaCl_2$ only treated cells, (* = $P < 0.05$, ** = $P < 0.01$, *** = $P < 0.001$, **** = $P < 0.0001$; no asterisk = not statistically significant).

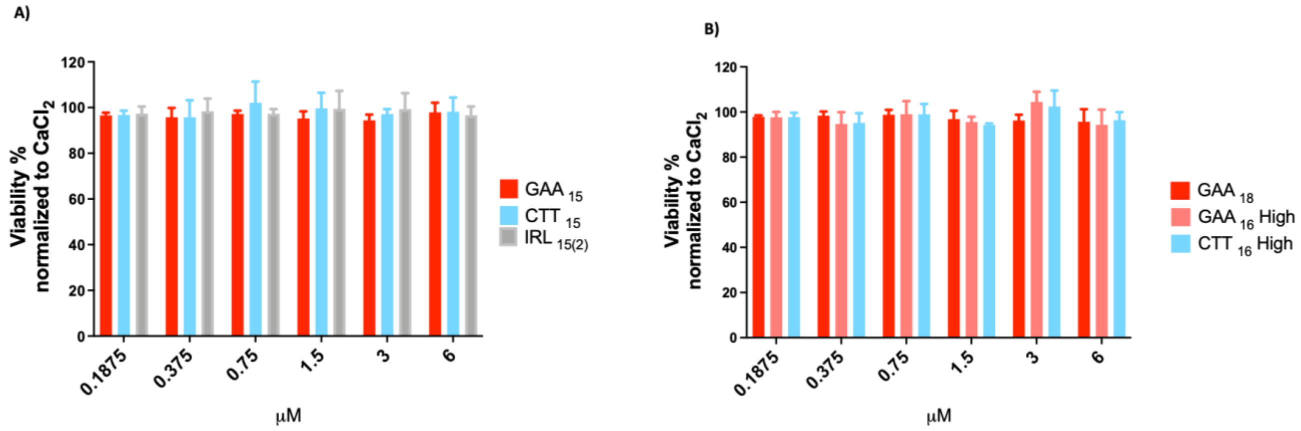


**Supplementary Figure 2.** No significant cytotoxicity was detected after treatment of FRDA derived patient's cells with selected ONs. Viability percentage of GM03816 cells after gymnotic delivery of ONs in medium supplemented with 9 mM $CaCl_2$ throughout 48 hours treatment. The relative values were obtained by normalization of ONs treated cells versus cells in the presence of 9 mM $CaCl_2$. Error bars = SD and (n ≥ 3).

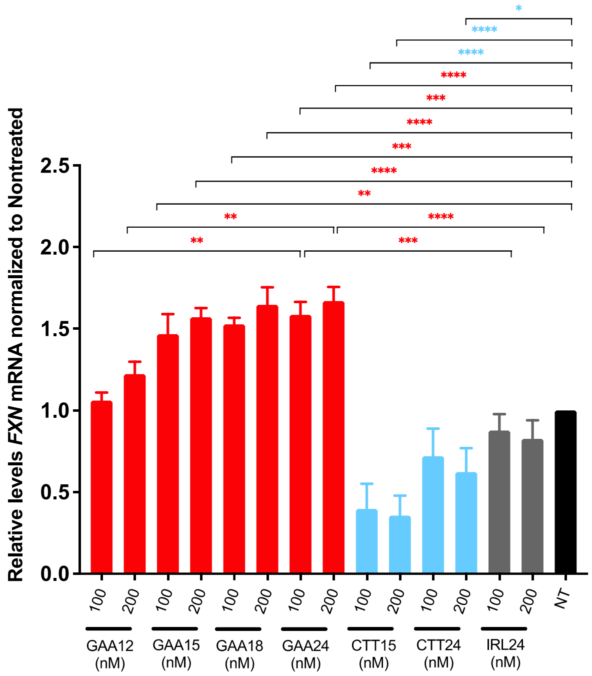


**Supplementary Figure 3.** GAA AGOs upregulate *FXN* mRNA expression in a FRDA cell model with a higher number of repeats. GM03665 fibroblasts were treated with different lengths of GAA, CTT and IRL ONs at 100 and 200 nM. Treated and nontreated cells were harvested 4 days after transfection, and *FXN* mRNA levels were assessed by RT-qPCR. The values were normalized to *HPRT* levels as reference gene and were compared to nontreated cells. Results are presented as Mean with SEM (n ≥3). Statistics were performed with two-way ANOVA Multiple Comparison, (Dunnett), towards nontreated cells. (* = $P < 0.05$, ** = $P < 0.01$, *** = $P < 0.001$, **** = $P < 0.0001$; no asterisk = not statistically significant). For both concentrations tested, $GAA_{15}$, $GAA_{18}$ and $GAA_{24}$ significantly upregulates *FXN* mRNA compared to nontreated cells. The treatment of $CTT_{15}$ at 100 and 200 nM and of $CTT_{24}$ at 200 nM resulted in significant downregulation of *FXN* mRNA.
